# A weighted sequence alignment strategy for gene structure annotation lift over from reference genome to a newly sequenced individual

**DOI:** 10.1101/615476

**Authors:** Baoxing Song, Qing Sang, Hai Wang, Huimin Pei, Fen Wang, XiangChao Gan

## Abstract

Genome sequences and gene structure annotation are very important for genomic analysis, while only the reference gene structure annotation is widely used for a wide range of investigations of different natural variation individuals. Herein, we are reporting the software GEAN which could lift over the reference gene structure annotation to other individuals belonging to the same or closely related species whose genome sequence was determined by whole-genome resequencing or *de novo* assembly. We found that inconsistent sequence alignment makes the coordinate lift over between different individual genomes unreliable, thus obscuring the lift over of gene structure annotations and genomic variants functional prediction. We designed a zebraic dynamic programming (ZDP) algorithm by providing different weights to different genetic features to refine the gene structure lift over. Using the lift over gene structure annotation as anchors, a base-pair resolution whole-genome-wide sequence alignment and variant calling pipeline for *de novo* assembly have been implemented. Taking *Arabidopsis thaliana* as example, we show that the natural variation alleles expression level of apoptosis death and defence response related genes might could be better quantified using GEAN. And GEAN could be used to refine the functional annotation of genetic variants, annotate *de novo* assembly genome sequence, detect syntenic blocks, improve the quantification of gene expression levels using RNA-seq data and genomic variants encoding for population genetic analysis. We expect that GEAN will be a standard gene structure annotation lift over and genome sequence alignment tool for the coming age of *de novo* assembly population genetics analysis.

## Introduction

Genome sequence and gene structure annotation are required for modern biological genome research. For large-scale population genetics projects, a high-quality reference genome sequence is usually assembled, and great efforts are exerted to generate the reference gene structure annotation. The genotypic data of a group of taxa are compared with the reference genome sequence. For whole-genome-wide resequencing variant calling projects, single-nucleotide polymorphisms (SNPs), insertions and deletions (INDELs), and other complex variants are generally represented by chromosome, coordinate, reference allele, and alternative allele (Danecek et al., 2011; Gan et al., 2011). While only the gene structure annotation of the reference sequence is generally used for the down-stream analysis e.g. gene expression level using RNA-seq, chip-seq. With the cost reduction of long-read sequencing technology and seeping up of genome assembly (Xiao et al., 2017; Ruan and Li, 2019), increasing natural variation individuals and species is being and will be sequenced with long reads and assembled *de novo* to reveal large-scale structure and rearrangements (Jain et al., 2018; Sun et al., 2018). Gene structure annotations of newly sequenced individuals are very important for many modern analyses, e.g., gene expression quantification, natural variation, population genetics, and strain-specific genes (Zhao et al., 2018; Lilue et al., 2018). While gene structure annotations of *de novo* assemblies have been predicated *ab initio*, and orthologous genes are detected by the alignment of only genetic sequences, which is independent of the genome sequence alignment (Zapata et al., 2016; Gan et al., 2016; Lilue et al., 2018). Algorithms have been developed to lift over the coordinates between different assembly versions (Hinrichs et al., 2006, 200; Spudich and Fernández-Suárez, 2010; Zhao et al., 2014). Here, we introduced the software GEAN, which could project the reference gene structure annotation to new genome sequence assemblies of the same or closely related species. Using the gene structure annotation as anchors, pipelines to perform base-pair resolution, whole-genome-wide pairwise and multiple sequence alignment (MSA), and variant calling has been implemented.

The lift over of the reference genomic coordinates to the pseudo-genome sequences of re-sequenced individuals could be performed with the variant calling records (Wang et al., 2010; MacArthur et al., 2012) by counting the number of base pairs shifted by the upstream variants. For the *de novo* assembly genomes, GEAN embeds an algorithm, which provides base-pair resolution, long region sequence alignment, and the reference coordinate could be lifted over with the refined sequence alignment. While the same variants might be represented in multiple ways, left alignment and whole-genome-wide multiple sequence alignment have been proposed to solve this problem for population genetics aim (Tan et al., 2015; Song et al., 2018). While the ambiguity of variant representation could cause false-positive open-reading-frame (ORF)-state interruption prediction (Fig. 1). And, there is rare well-designed software to perform genome sequence alignment and lift over the reference gene structure annotation to a new assembly that show considerations for gene structure. Thus, the generation of false-positive ORF-shift predication and improper genotypic variant functional predication is expected.

**Fig. 1.**
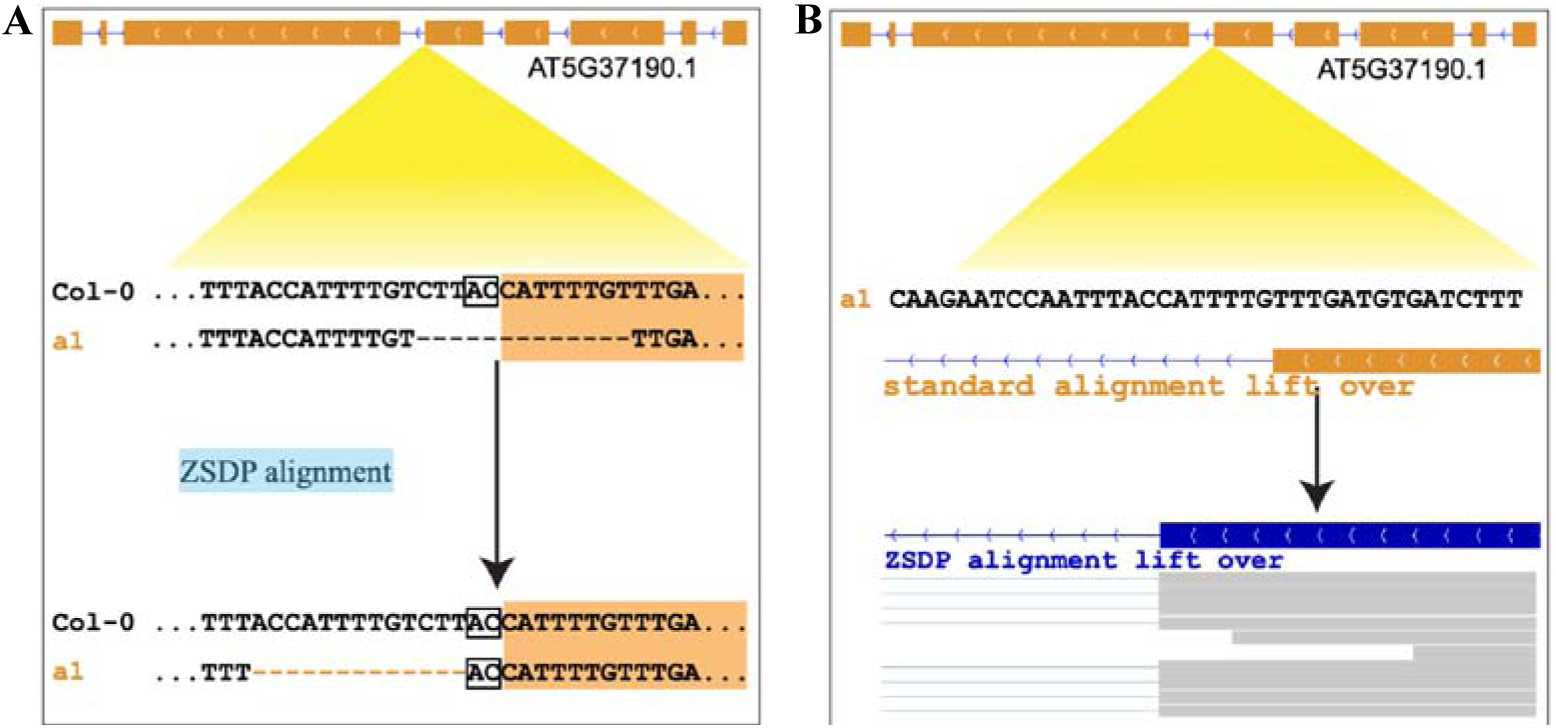
Example of inconsistent sequence alignments that affect variants functional inference. (A) Upper panel standard sequence alignment suggests that a 13 bp deletion disturbs the splice site and shifts the ORF of AT5G37190.1. Lower panel alignment, updated using ZSDP, suggests that a 13 bp deletion is located in the intron, and the splice site is conserved. (B) Upper panel shows the gene structure annotation of AT5G37190.1 of a1 allele by coordinate lift over using standard sequence alignment. Lower panel shows the gene structure annotation updated by the ZSDP algorithm, and the RNA-seq reads mapping support the lower panel.

Here, we provide a semi-global sequence alignment algorithm and software to infer about the gene structure of non-reference accession/line with an algorithm called zebraic dynamic programming (ZDP). Dynamic programming could be accelerated by extending the striped Smith-Waterman (SSW) algorithm (Farrar, 2007), and the accelerated version of ZDP is called zebraic striped dynamic programming (ZSDP). We demonstrate that the ZSDP algorithm result is supported by RNA-seq data. And this method helps to infer the functional impact of variants with the data from *Arabidopsis thaliana* 1001 Genomes project (Alonso-Blanco et al., 2016), gene expression level quantification, annotate *de novo* assembly genome sequence, and perform variant calling for the *de novo* assembly of the genome sequence (Zapata et al., 2016; Gan et al., 2016; Jiao et al., 2017; Springer et al., 2018). Moreover, we are extending this method to other genomic fields.

In the subsequent sections, we demonstrate the performance of GEAN by using *Arabidopsis thaliana*, *Cardamine hirsuta* (Gan et al., 2016), and *Drosophila melanogaster* genome sequences and related evidence. First, by using the *Arabidopsis thaliana* and *Drosophila melanogaster* genomes, we develop concepts that underlie the algorithm of GEAN as a tool to provide gene structure annotation for different inner species individuals. We then consider the inter-species genomes and try to transform the gene structure annotation of *Arabidopsis thaliana* to *Cardamine hirsuta*. We also show that the lift over gene structure annotation could help perform variant calling for the *de novo* assembly genome sequence and whole-genome-wide multiple sequence alignment (MSA).

### Lift over of genomic coordinates

Advances in and decreasing costs of high-throughput sequencing have revolutionized the ability to investigate sequence diversity, thereby allowing the international consortium of geneticists to develop foundational resources (Consortium, 2012; Consortium, 2015; Alonso-Blanco et al., 2016, 1; Wang et al., 2018). The compatibility and the ability to integrate different accessions become very interesting. One of the crucial steps is to lift over the reference coordinates, under investigation, to another genome sequence, so that we can compare the orthologous haplotype sequences of interesting regions from two or multiple different individuals.

For the whole-genome resequencing data, the pseudo genome sequence could be obtained by replacing the reference alleles with alternative alleles taking vcf (Danecek et al., 2011) or sdi (Gan et al., 2011) file as input. We embed an algorithm in GEAN to generate the pseudo genome sequence from variant calling result and lift over reference coordinates to the pseudo genome. The lift over of the gene structure could be performed by the lift over of each boundary coordinate.

To lift over the reference coordinates to another *de novo* assembly genome sequence, initial sequence alignment could be performed using the available low-resolution methods for whole genome sequence (Marçais et al., 2018; Li, 2018) or genetic sequence (Li, 2018). Taking low-resolution genome alignment as anchors, GEAN uses a sliding window (Methods) to perform base-pair resolution (Fig. 2), whole-genome wide sequence alignment and lift over the gene structure annotation with the refined sequence alignment.

**Fig. 2.**
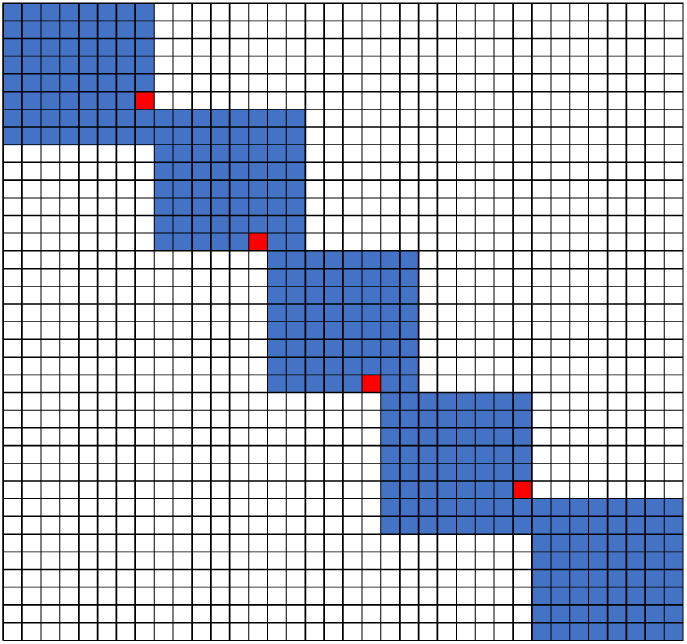
Sliding window sequence alignment method implemented in GEAN for the long sequence alignment (Methods).

### Mitigation of false deleterious variants caused by inconsistent alignment

A INDEL/SNP ratio of ∼25% has been observed in the population genomes of plant, animal and human beings (Rakocevic et al., 2019), and at least 63%-65% INDEL might be affected by inconsistent alignment in a population (Song et al., 2018). As shown in Fig. 1A, the standard sequence alignment based gene structure projection could lead to gene false loss-of-function prediction and improper variants functional prediction. The genetic load of genotypical variants has been analyzed for various purposes by using the data from whole-genome resequencing projects, e.g., GWAS burden test (Song et al., 2018), deleterious mutations (Ramu et al., 2017) and non-synonymous/synonymous mutation ratio for natural selection analysis (Nekrutenko et al., 2002; Liu et al., 2008). Here, we show the number of transcripts, without variants shifting ORF and disturbing splicing sites, could be falsely predicted as loss-of-function.

By using the variant calling result of 1, 211 *Arabidopsis thaliana* accessions (Alonso-Blanco et al., 2016) and 203 *Drosophila melanogaster* lines (Huang et al., 2014; Dembeck et al., 2015) with IMR/DENOM (Gan et al., 2011), we created the pseudo genome sequence of each accession and performed gene structure lift over with simple coordinate lift over firstly. For the lifted protein coding transcripts, which are predicted as loss-of-function, we realigned the structure annotation by using the ZSDP algorithm (Fig. 3). A total of 6,713 transcripts have been realigned as ORF state conserved for *Arabidopsis thaliana* accessions and 1,946 transcripts for *Drosophila melanogaster*. An average of 158 and 47 transcripts have been re-aligned as ORF state conserved for each *Arabidopsis thaliana* and *Drosophila melanogaster* accession, respectively. We observed negative correlations (p value < 2.2e-16) between the number of realigned transcripts and identity by state (IBS) index for both species (Fig. 4), which suggested that more transcripts have been affected by inconsistent alignment between individuals with increased diversity.

**Fig. 3.**
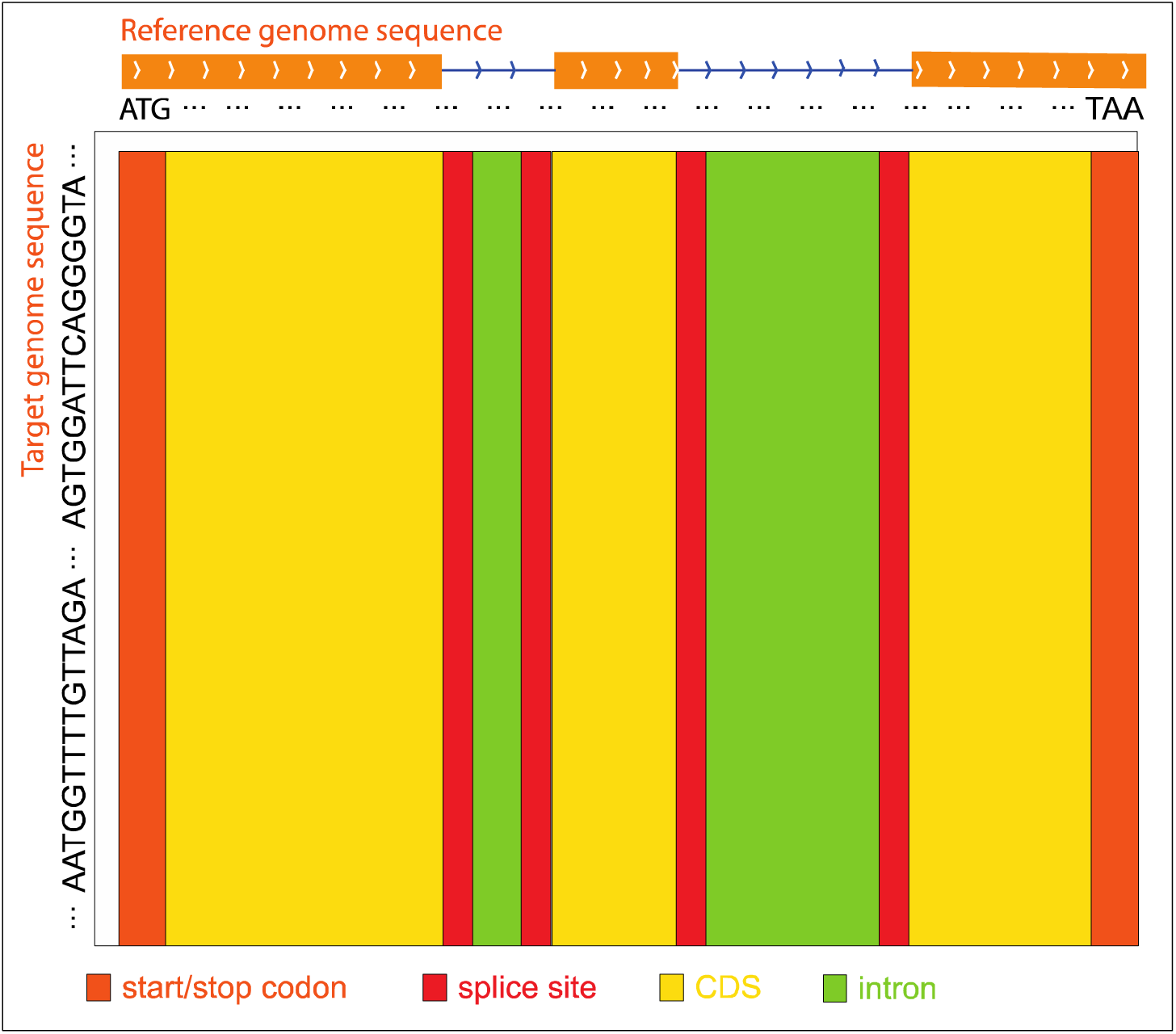
ZSDP sequence alignment methods. When the reference genome sequence is aligned to the target accession genome sequence, different score strategies are used for different reference genetics region to construct the dynamic programming score matrix. The purpose is to align the CDS regions preferentially.

**Fig. 4.**
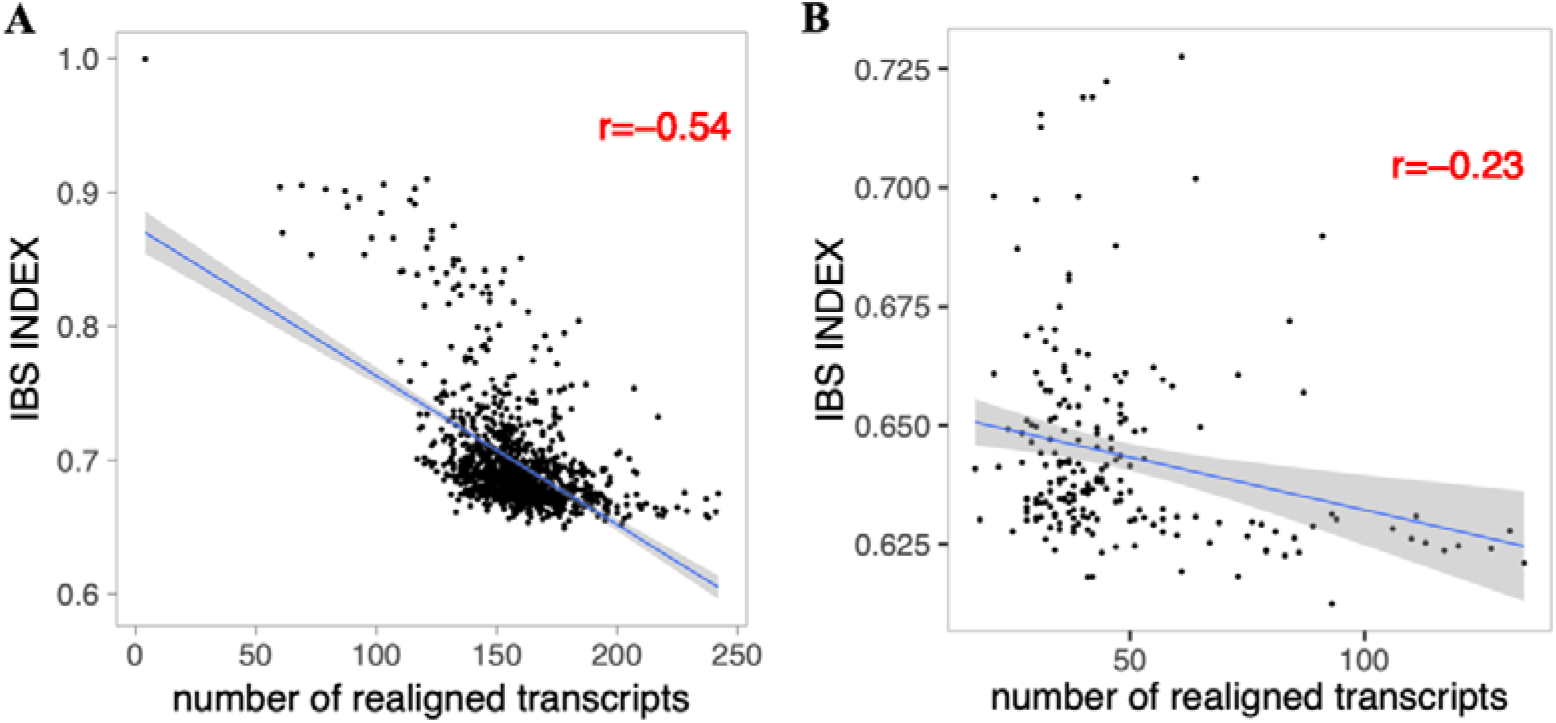
Correlation between the number of realigned transcripts by the ZSDP method and IBS index for *Arabidopsis thaliana* (A) and *Drosophila melanogaster* (B).

To show the advantage the ZSDP approach, we compared the created gene structure annotation file with the RNA-seq data using Cufflinks v2.2.1 (Trapnell et al., 2012). Raw RNA-seq reads of 728 *Arabidopsis thaliana* accessions released by the *Arabidopsis* 1001 Epigenomes project (Kawakatsu et al., 2016), were and mapped to pseudo genome sequence. After filtering low-quality accession (less than 3,000,000 RNA-seq reads available, alignment rate ≤90%), 697 accessions were left. And the generated bam files were feed into Cufflinks v2.2.1 to get all the splice site positions. For all the controversial splice sites between simple coordinate lift over and ZSDP, 41,235 controversial split sites could not be determined by the RNA-seq reads due to the absence or low expression level, and the RNA-seq data confirmed 23,448 ZSDP alignment splice sites, which is ∼20 times that of the standard sequence alignment lift over of splice sites (1,200).

For regions with potential false loss-of-function predication variant records, GEAN realigns the transcripts, recall the variants, and replace the old variants with the re-aligned variant records. The newly re-aligned variant records could be used to predict the functional annotation of genetic variants (Wang et al., 2010).

### Non-reference line RNA-seq reads mapping rate could be improved using pseudo genome sequence

For many RNA-seq projects, regardless of what materials is used, the reference genome sequence is usually used for RNA-seq read mapping (Krizek et al., 2016; Kawakatsu et al., 2016). Here, we tested whether mapping read to the pseudo genome sequence could generate an increased mapping rate and if there any gene show different expression level between those two approaches. For this aim we set up a pipeline to generate a compressive gene structure annotation of pseudo genome sequence. Using GFF file from coordinate lift over, the ORF states of the target line could be checked with the inferenced haplotype sequence. Any ORF-state disruption according to the lift over results will be realigned with the ZSDP algorithm.

The ZSDP method could migrate the reference gene structure annotation to the pseudo genome sequence. However, the new accession maybe has some gene structure variation i.e. number of exons that changed, alternative splice site, novel gene. So in GEAN we complemented the gene structure lift over with orthologue-based gene structure annotation (Slater and Birney, 2005) and the results of external annotation method (e.g., *ab initio* annotation (Stanke and Waack, 2003) and transcript assembly (Trapnell et al., 2012)). As illustrated in Fig. S1, the gene structure predicated by the upstream module would be adapted firstly, and the region predicted as ORF-state shift or non-coding would be handled by the below modules. An average of 346 transcripts was re-annotated by using ZSDP or Exonerate (Slater and Birney, 2005) for each accession.

We mapped RNA-seq reads of 728 *Arabidopsis thaliana* accessions (Kawakatsu et al., 2016) to Col-0 reference genome sequence and pseudo genome sequence separately. The gene structure annotations for each pseudo genome were obtained using the whole gene structure annotation pipeline in GEAN. *Ab initio* gene structure predications were used for the fifth modules of our pipeline. Raw RNA-seq were trimmed and mapped as descripted above. Mapped reads were counted using HTSeq 0.9.1 (Anders et al., 2015) with default parameters. The mapping rate to the pseudo genome sequence was significantly higher than that to Col-0 reference genome sequence (94.76% vs. 96.33, p value < 2.2e-16, Wilcoxon signed rank test) (Fig. 5).

**Fig. 5.**
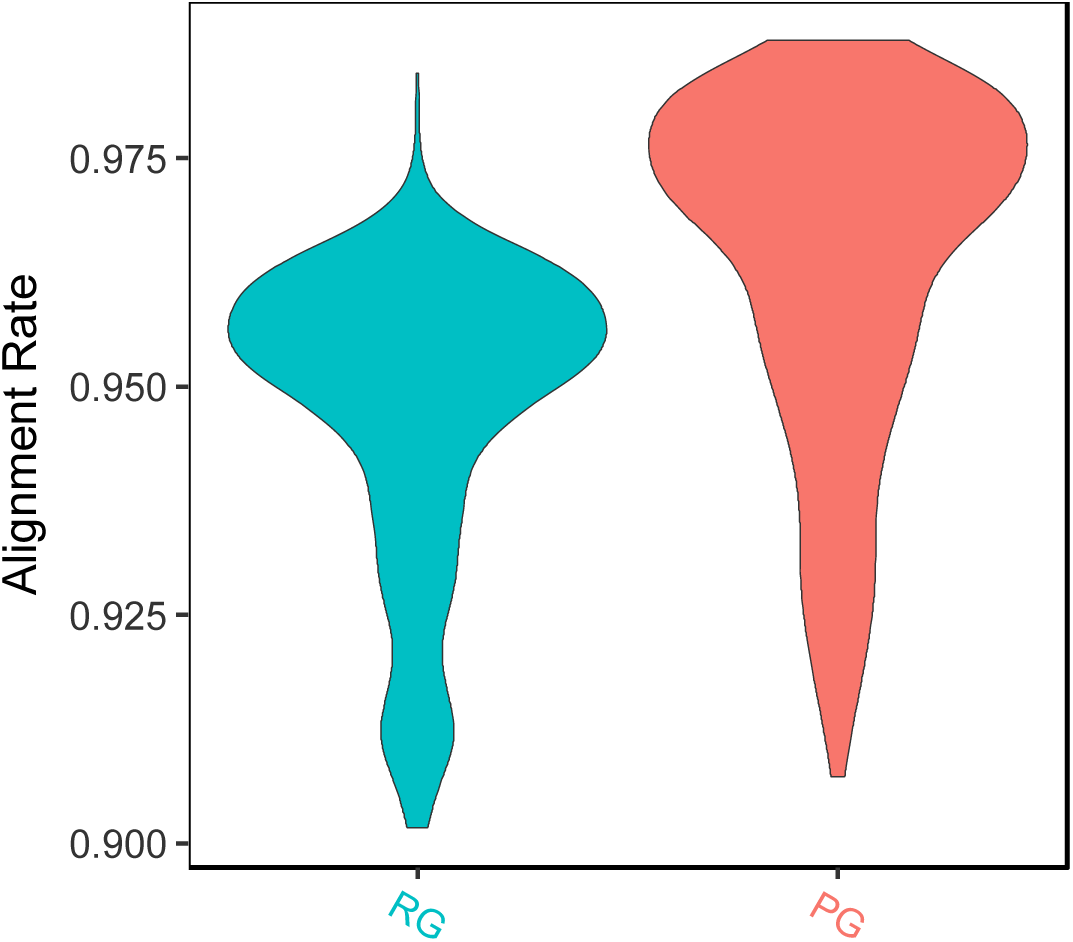
Mapping rate of RNA-seq read to pseudo genome (PG) sequence and Col-0 reference genome (RG) sequence.

Due to improved mapping rate, a group of 5,122 genes had significantly higher expression level when quantified by pseudo genome gene structure annotation. And the sequence diversity (measured using pi value) of these genes is significantly higher than that of the whole genome wide background (Wilcoxon test p value < 2.2e-16). GO analysis was conducted using agriGO (Du et al., 2010) with default settings. These results suggested that the advantage of GEAN pipeline for gene expression level quantification is essential for more diverse genes (Fig. 6), especially genes related to cell death, defense response, and immune response (Fig. 7).

**Fig. 6.**
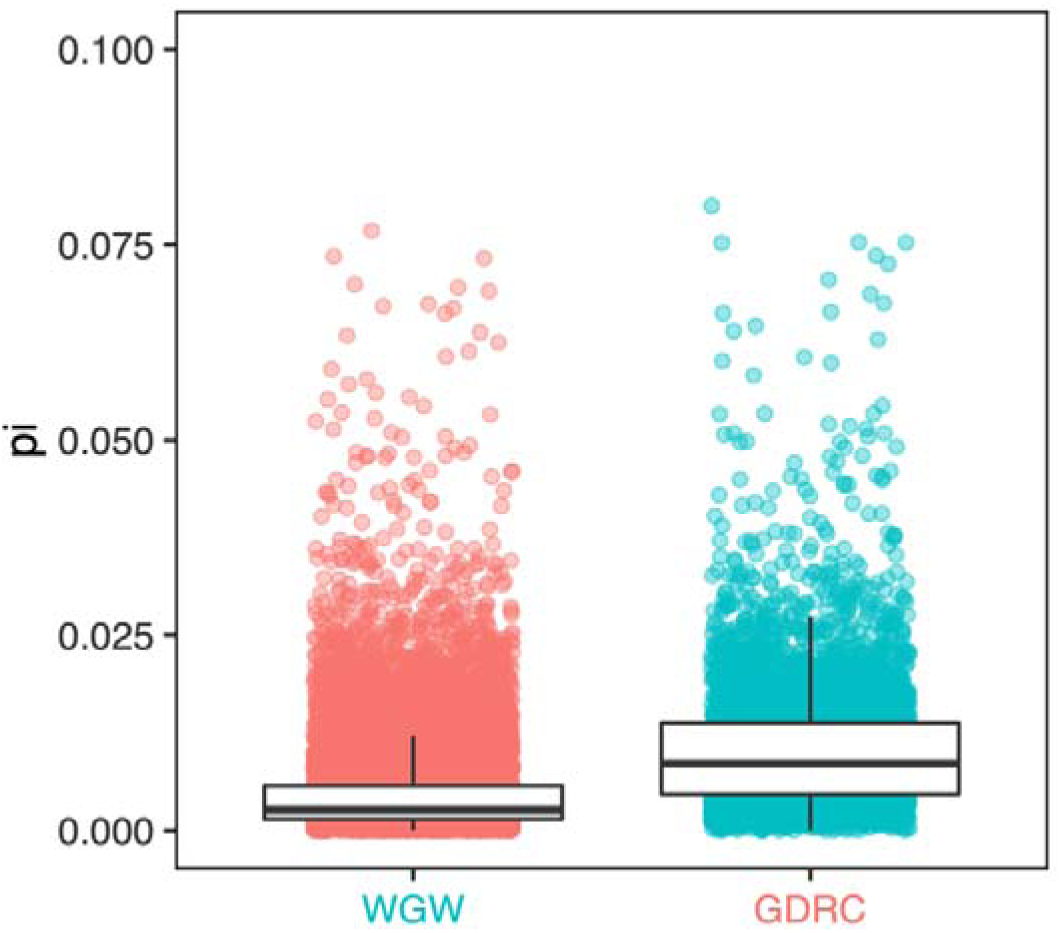
Sequence diversity of genes with significantly different read counts (GDRC) versus the whole-genome wide (WGW) background.

**Fig. 7.**
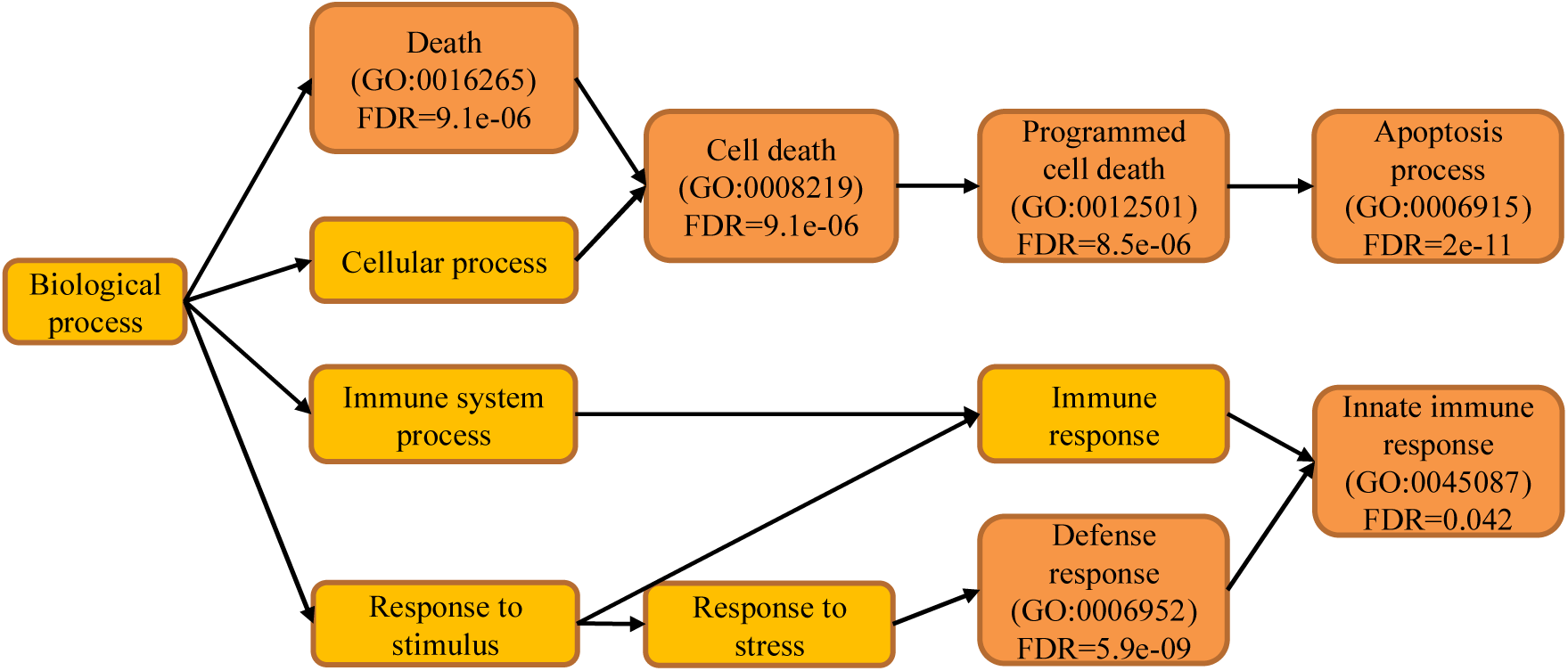
Directed acyclic graph of GO biological process terms that are enriched (dark orange) in a set of 5,122 genes showing different expression levels in at least 5 out of 697 accessions in which the RNA-seq reads mapping rate was greater than 90 using both Col-0 genome sequence and pseudo genome as reference.

### Transformation of the reference gene structure annotation to *de novo* assembly genome sequence

The comparison of the genomic variants of different species and individuals is a very interesting topic. With the reduced cost of long-read sequencing technologies, the genomes of many species have been or will be sequenced. Some inner-species population sequencing projects are moving from short-read to long-read technology. For comparative genomic analysis, aligning the *de novo* assembly to reference accession or well-developed model organisms is an important step. The genome alignment information could be used to transform the reference gene structure annotation to the newly assembled genome sequence.

We implemented a gene structure annotation function in GEAN by utilizing the genome sequence alignment information. The genome alignment result is usually denoted by a pair of ranges: one is from the reference sequence and the other is from the query genome sequence. ZSDP performs pairwise sequence alignment with a very efficient sliding window method. It lifts the reference gene structure annotation to the query genome sequence and updates the annotation using the ZSDP algorithm.

We tried to transform the gene structure annotation of *Arabidopsis thaliana* Col-0 to *Arabidopsis thaliana* L*er*-0 and *Cardamine hirsuta* reference genome sequence. *Arabidopsis thaliana* L*er*-0 was assembled into the chromosome level (Zapata et al., 2016). *Cardamine hirsuta* is a species closely related to *Arabidopsis thaliana* phylogenetically, whose genome has been published (Gan et al., 2016). We found that 34,869 out of 39087 protein coding transcripts of *Arabidopsis thaliana* Col-0 accession could be transformed to L*er*-0, 32,307 transformed annotations had a conserved ORF state, and 752 of them were re-aligned by the ZSDP algorithm. A total of 14,905 Col-0 protein coding transcripts could be transformed to *Cardamine hirsuta* as conserved ORF-state, while 7,119 of these were re-aligned by the ZSDP algorithm. The higher proportion of transcripts realigned by ZSDP in *Cardamine hirsuta* was confirmed based on the negative correlation between the number of realigned transcripts and IBS index. A dot plot of the transformed gene annotation from Col-0 to *Cardamine hirsuta* was comparable with the Circos plot, showing synteny between the genomes of *Arabidopsis thaliana* and *Cardamine hirsuta* published before (Gan et al., 2016). In addition, a large proportion of genes that did not follow the large synteny fragments were observed in *Cardamine hirsuta* than in L*er*-0. Several syntenic analysis modules have been implemented in the GEAN software taking the reference gene structure annotation file and the lift over gene structure annotation file as input (Fig. 8).

**Fig. 8.**
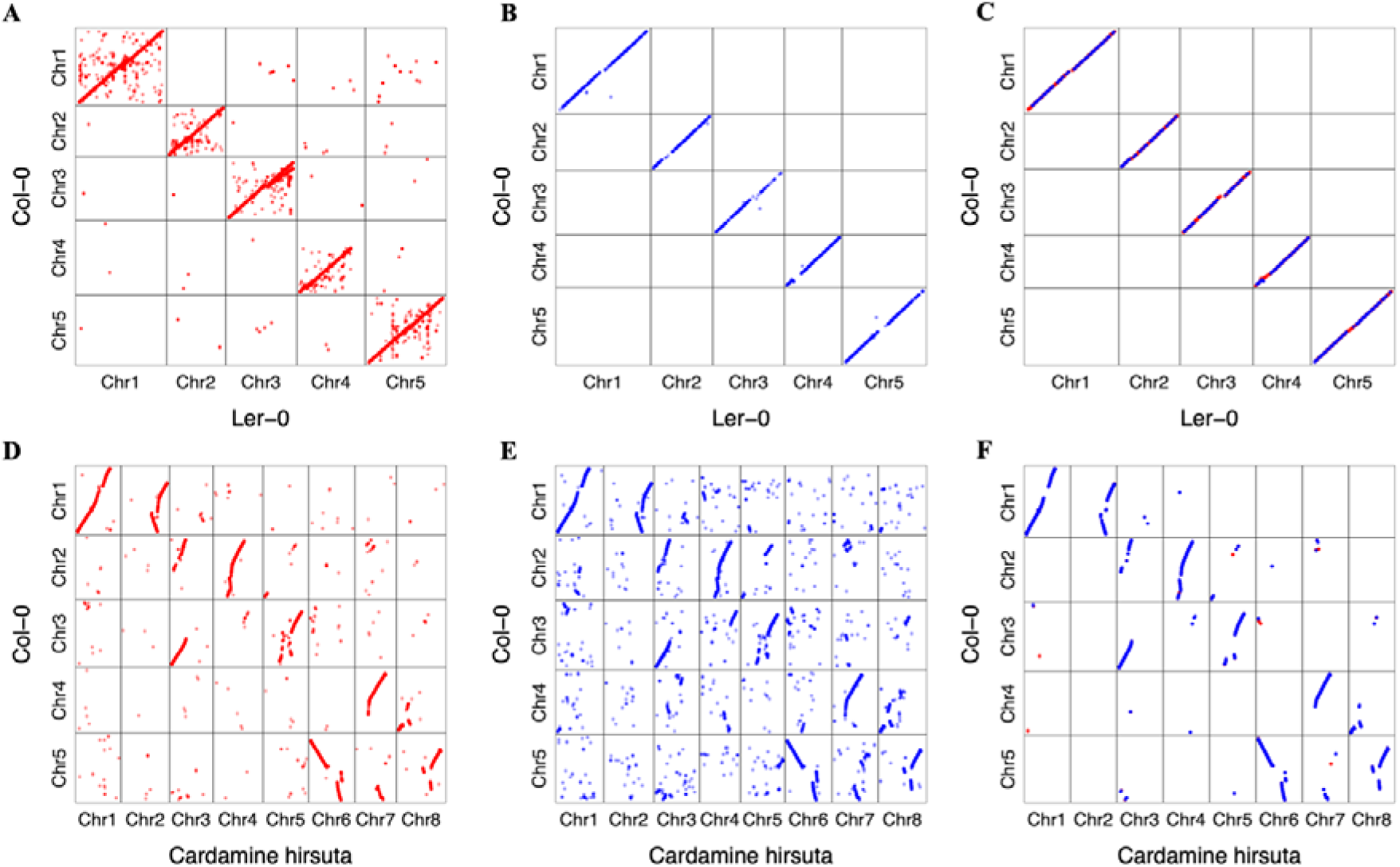
Dot plot of gene position in Col-0 against the position of genes transformed to other *de novo* assemblies. Dots in red color are the result from standard sequence alignment lift over directly, and those in blue color have been realigned by ZSDP algorithm. (A) transform *Arabidopsis thaliana* Col-0 gene structure annotation to L*er*-0 accession genome sequence using standard sequence alignment. (B) transform Col-0 gene structure annotation to L*er*-0 genome sequence of those genes could not be transformed using standard sequence alignment using ZSDP algorithm. (C) investigate syntenic blocks between Col-0 and L*er*-0 using the transformed gene structure annotations. (D) transform *Arabidopsis thaliana* Col-0 gene structure annotation to *Cardamine hirsuta* genome sequence using standard sequence alignment. (E) transform Col-0 gene structure annotation to *Cardamine hirsuta* genome sequence of those genes could not be transformed using standard sequence alignment using ZSDP algorithm. (F) investigate syntenic blocks between *Arabidopsis thaliana* Col-0 and *Cardamine hirsuta* using the transformed gene structure annotations.

For a comprehensive annotation of *de novo* assembly genome sequence, the GEAN results could be used as a very reliable information source for a complete gene annotation pipeline (Haas et al., 2008).

### Base-pair resolution variant calling for *de novo* assembly genome sequence

Variant calling is important for various population genomics analysis. We implemented a pipeline in GEAN to perform whole-genome wide sequence alignment, and variant calling using the lift over gene structure annotation as anchors. In detail, we aligned the gene structure annotation of *de novo* assembly genome sequence with the reference gene structure annotation by using a Needleman-Wunsch algorithm and treating identical gene ID as match. The start and stop codons of matched genes were used as anchors to split the whole genome sequence into fragments, and the sliding window method was used to perform base-pair resolution sequence alignment and variant calling in each fragment. By applying this method on a L*er*-0 *de novo* assembly genome sequence (Zapata et al., 2016), 2,664,186 SNPs and 546,778 INDELs were detected, which are higher than those from whole genome resequencing variant calling (693,834 SNPs and 159,350 INDELs) (Song et al., 2018). Except for the probability that *de novo* assembly has a high power of variant calling in highly diverse regions, we classified all the variants into SNP and INDEL only. Hence, for the structure variation, e.g. relocation, could produce a cluster of variant records. Interestingly, the INDEL/SNP ratios for the two results are comparable with each other. Any two of the reference genome sequence, the variant calling result and *de novo* assembly are enough to inference the third one. As far as we know, GEAN is the first software could call variant in such a comprehensive manner.

### Whole-genome wide MSA

For the variant calling of a population of individuals, inconsistent alignments could generate inconsistent variant records. Previously, we showed that this problem could be solved with whole MSA (Song et al., 2018). While, both the left alignment method (Tan et al., 2015) and genome-wide MSA did not consider the gene structure.

GEAN performs MSA for each genetic feature separately. To make is faster, GEAN try to use short and non-overlapping fragments to run the MSA algorithm. Considering that the boundary of genetic features of functional conserved transcripts could be well defined, GEAN performs MSA for short elements (e.g. intron, CDS, intergenic region) directly and only uses an overlapping sliding window to perform MSA only for long elements.

## Method

### Definitions

The ORF state has defined in the same way as (Song et al., 2018).

### Coordination lift over

Considering the presence of INDELs, SVs, and even sequence fragment rearrangements, the coordinates of a certain orthologous haplotype sequence fragments are different from each individual. Lift over is a way of mapping coordinate from one genome assembly to another.

For whole-genome resequencing projects, GEAN performs lift over by counting the number of base pairs that were shifted by the upstream variants. For the *de novo* assembly sequence, considering the similar range entries as input (e.g., Mummer(Marçais et al., 2018), minimap2(Li, 2018, 2)), GEAN generates the base-pair resolution pairwise sequence alignment with a dynamic programming algorithm by using a sliding window, which would be used for the coordinate lift over.

### Zebraic dynamic programming

We designed a pairwise sequence alignment algorithm taking gene structure into consideration by extending the available dynamic programming algorithms (Needleman and Wunsch, 1970; Smith and Waterman, 1981; Gotoh, 1982). The two sequences to be compared, query sequence and reference sequence, are defined as Q=q_1_, q_2_ … q_m_ and D=d_1_, d_2_ … d_2_, respectively. The length of the query and database sequences are m=|Q| and n=|D|, respectively. A scoring matrix W(q_i_, d_j_) is defined for all residue pairs. The score W(q_i_, d_j_) ≤0 when q_i_≠d_j_, and W(q_i_, d_j_) > 0 when q_i_=d_j_. The penalty for starting and extending a gap are defined as G_init_ and G_ext_, respectively.

As we know the gene structure of the reference sequence, i.e., intron, CDS, start codon, stop codon, and splice sites. So, we use the reference sequence from start codon to stop codon. And the query sequence is in general extend with variant calling based coordinate lift over, both upstream and downstream by a length of the gene length, to make sure the genetic region in include in the selected region. When constructing the score matrix, we initialize the first row and the first column with 0. At the traceback step, we start the tracing from the cell with the largest value in the last column, since we expect the reference sequence could be globally aligned while the query sequence could be locally aligned. And different W(q_i_, d_j_), G_init_, and G_ext_ values were used for intron, CDS, splice sites, and start and stop codons separately, so we have all the score strategies as W(q_i_, d_j_)_intron_, G_init_intron_, G_ext_intron_, W(q_i_, dj)CDS, Ginit_ CDS, Gext_CDS, W(qi, dj)spliceSites, Ginit_ spliceSites, Gext_ spliceSites, W(qi, dj)start/stopCodon, G_init_ start/stopCodon_, and G_ext_ start/stopCodon_. The protein coding region and splice sites will be highly weighted, and the sequence in those regions will be aligned primarily. Thus, any ambiguity in ORF-states caused by different genetic variance representation could be polished to keep the ORF-states complete.

As we need to determine which gene structure elements of the current j are located and avoid repeating this query process for scoring matrix filling and traceback, we defined a tracking matrix T with the same size as the score matrix, which records the scoring path of each cell of the score matrix. The value of each cell in the track matrix is a certain value SNP, INSERTION, DELETION, or an ambiguous value SNPORINSERTION, SNPORDELETION, INSERTIONORDELETION, SNPORINSERTIONORDELETION.

The alignment scores ending with a gap along D and Q are E calculated using Equation (1) and F calculated using Equation (2), respectively:

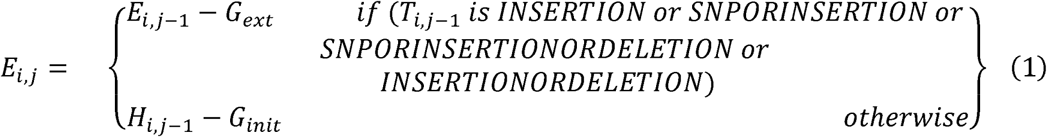

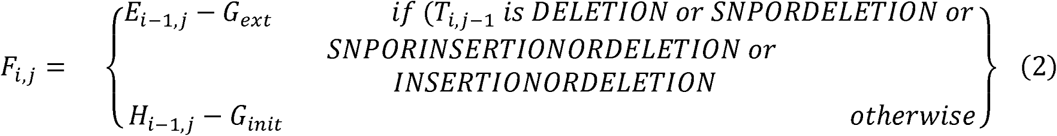

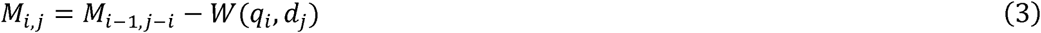

The alignment score for H_i,j_ where 1≤i≤m and 1≤j≤n is defined by Equation (4):

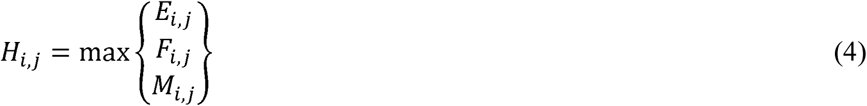

The matrix T was updated as:

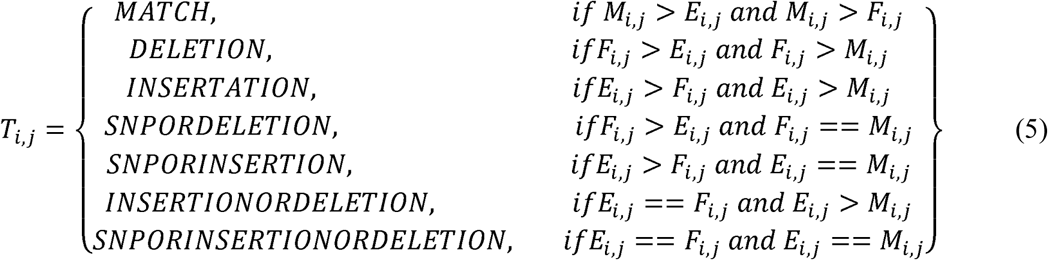

Once, *T_i,j_* is set as a certain value i.e. SNP, INSERTION, DELETION, SNPORINSERTION, all the continuous ambiguous ancestral cells will be set as certain value, until a certain value being found. For RAM-saving purposes, instead of keeping the whole H_i,j_ matrix, GEAN only keeps the last row and last column, and the traceback step can be easily performed using the tracking matrix.

### ZSDP for gene structure alignment

ZSDP is a speeding up version of ZDP algorithm by intra-sequence parallelization, which parallelize the algorithm by extending the SSW algorithm (Farrar, 2007) to a semi-global sequence alignment manner. The SSW algorithm could be 10 times faster than the standard smith-waterman algorithm using AVX2 and has been widely embedded in several high-throughput sequencing read mapping software, such as Bowtie2 (Langmead and Salzberg, 2012, 2), BWA-SW (Li and Durbin, 2010), and Stampy (Lunter and Goodson, 2011).

In the SSW algorithm, when calculating H_i,j_, the value from the scoring matrix W(q_i_, d_j_) is added to H_i-1, j-1_. To avoid looking up for W(q_i,j_) for each cell, a query profile parallel to the query is calculated for each possible residue. The query profile is calculated once for each database search. The calculation of H_i,j_ only requires the addition of the pre-calculated score to the previous H_i,j_. The ZSDP method takes a similar approach by pre-calculating a query profile for each gene structure element category separately, and uses a standard method to implement the lazy F evaluation loop.

The SSW method only provides the optimal alignment score but does not report the information necessary to construct the final alignment. The SSW Library reports the detailed alignment by performing SSW twice (get the ending positions by a forward SSW and then generate the beginning position by a backward SSW) and performs a standard smith-waterman sequences alignment between the beginning and ending position (Zhao et al., 2013), which should be further optimized.

Here, we constructed the H_i,j_ vector by comparing H_i-1,j-1_ and E_i,j_ and stored the H_i,j_ from vHstore into the standard integer scoring matrix. We calculated the F_i,j_ score for each cell of the current column. By comparing the F_i,j_ value with the H_i,j_ value, we updated the H_i,j_ and vHstore with the larger value. The corresponding value in the traceback matrix was also updated. Then, the traceback step of the dynamic programming algorithm, which provides the detailed alignment, could be performed using the traceback matrix.

The computational speeding up is related to the number of cells calculated per CPU register. We used 8, 16, and 32 bits to process the score matrix for sequence with different lengths. So that we could place as many cells as possible into each single instruction multiple data register. Meanwhile, the integer type is wide enough to process the sequence alignment scores. We implemented the striped dynamic algorithm with AVX2. AVX2 is available for most of model Intel processors, whose register is 256 bits wide.

### Sequence alignment by using the sliding window

To accelerate the alignment long sequence, we used a sliding window. For each window, we determined the maximum value of the last row or column. The maximum cell would be used as the start cell of the next scoring window. With this strategy, we have a linear computational time costing. Similar with banded alignment, we note that this type of heuristics can fail to give the optimize alignment under some situations. Luckily, the failures are always only present in small local regions. A large sliding window size could be used to avoid this problem, which is parameter could be passed from command line. This problem could be total avoided, if the sliding window is larger than the sequence being aligned.

### Gene expression level quantification

Raw RNA-seq reads were filtered with Trimmomatic (Bolger et al., 2014) (v0.36, LEADING:10 TRAILING:10 SLIDINGWINDOW:4:15 MINLEN:6) and mapped to genome sequence using Hisat2 (Kim et al., 2015) (v2.1.0, --max-intronlen 30000).

*Ab initio* gene structure predications were performed using Augustus (Stanke and Waack, 2003) (--genemodel=complete --maxDNAPieceSize=2000000 --sample=200).

Significantly higher expression qualified using reference genome sequence and using pseudo genome sequence are roughly defined as data out of range lower fence Q1-5*IQ and upper fence Q3+5*IQ, Q1 is the 25th percentile, Q3 is the 75th percentile, and IQ=Q3−Q1. And the analysis was performed using R command boxplot.stats(log(reads_count)ratio, coef=5)$count.

### Software

The gene structure annotation alignment is implemented as the GEAN software. Related functions, such as coordinate lift over any position, obtaining the pseudo genome sequence, and checking the ORF states, are also implemented. Additionally, a function that lifts over all the reference gene structure annotation to other accessions is implemented.

GEAN is written in C++ and parallelized using the C++ Standard Library and freely available at https://github.com/baoxingsong/GEAN. A series of examples and tests is embraced in the distribution. By making the system open source, we hope to encourage others to expand and improve the code base.

## Conclusions

As the examples above show, the capabilities of GEAN enable a researcher to transform the gene structure annotation of any two or collections of genomic sequences. This could be used for RNA-seq analysis and perform base-pair resolution, whole-genome-wide sequence alignment by using computer facilities that are widely available today.

## Declarations

### List of abbreviations

ZDP: zebraic dynamic programming
SNP: single-nucleotide polymorphisms
INDEL: insertions and deletions
MSA: multiple sequence alignment
ORF: open-reading-frame
SSW: striped Smith–Waterman
ZSDP: zebraic striped dynamic programming
IBS: identical by state

### Competing interests

The authors declare that they have no competing interests.

### Funding

This work was supported by Guizhou Science and Technology Department grants QKH Basic research [2019]1298 and Guizhou Provincial Education Department grants QJH-KY-Z [2018]420 to F.W.

## Authors’ contributions

BS, QS, FW and XG conceived this software. BS and QS structured the draft and provided final editing. BS coordinated and drafted the manuscript, and synthesized comments provided by all authors. All authors contributed critically important comments. BS implemented the software, HP and FW performed testing and contributed source code. All authors read and approved the final manuscript.

## Acknowledgments

We thank Edward S. Buckler and Miltos Tsiantis for the suggestions and comments on the work; Hequan Sun and Bjorn Pieper for discussions; Elad Oren for extending the usage to genome sequence *de novo* assembly quality control. We also acknowledge Sara Miller for the revision.

## Authors’ information

BS is working as a postdoc at Ed Buckler group at Cornell University. Qing Sang is a postdoc at George Coupland group Max Planck Institute for Plant Breeding Research. FW and HP are working as a lecturer in Qiannan Normal College for Nationalities. XG is a group leader at Max Planck Institute for Plant Breeding Research.

## Supplementary information

**Fig. S1.**
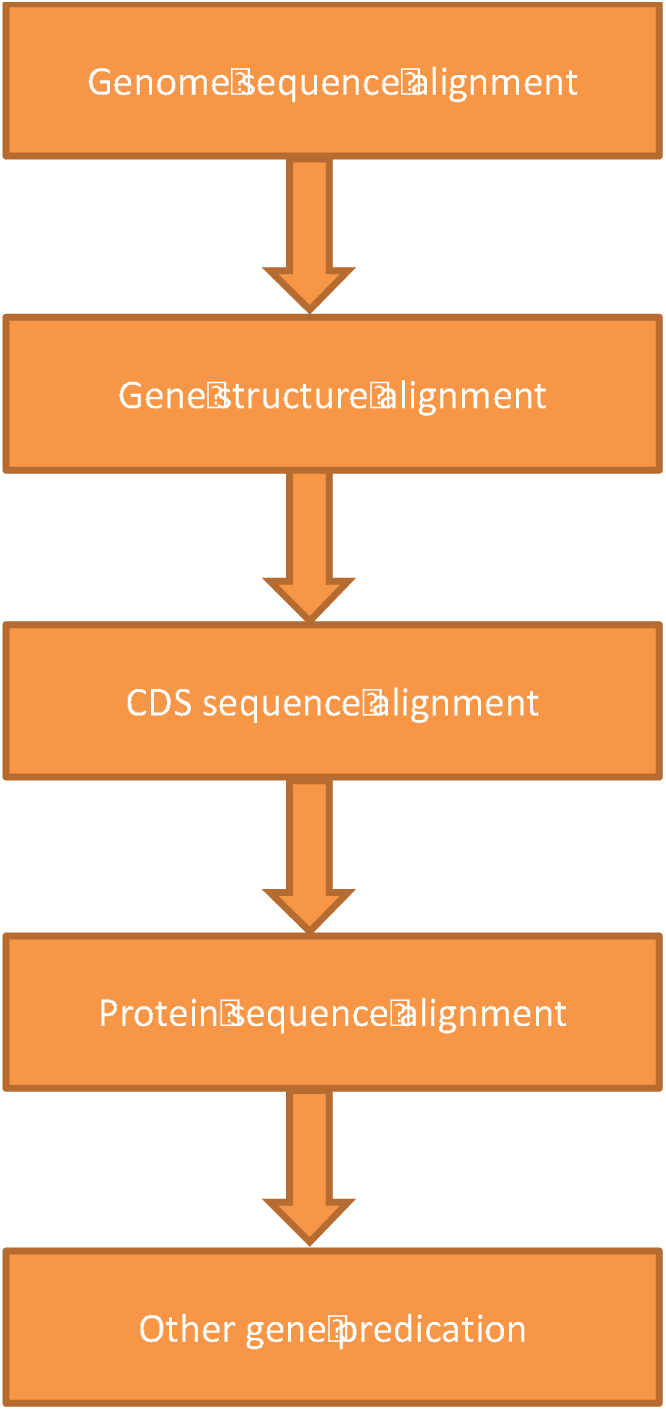
Schematic showing the pipeline for the protein coding gene annotation of the non-reference genome. The gene structure annotation of upstream step would be supplemented with the downstream method.

